# Woody plant encroachment enhances negative interactions in Arctic plant communities over the last 24,000 years

**DOI:** 10.1101/2025.05.26.656118

**Authors:** Ying Liu, Weihan Jia, Sisi Liu, Simeon Lisovski, Kathleen R. Stoof-Leichsenring, Luca Zsofia Farkas, Ulrike Herzschuh

## Abstract

Woody taxa encroachment in the Arctic has been widely observed. However, it remains uncertain how plant interactions are affected by such encroachment due to the lack of long-term observational data. Here, we reconstruct plant composition and functional trait turnover during post-glacial woody encroachment using sedimentary ancient DNA from nine lakes in the Siberia-Alaska region. Environmentally constrained null models are applied to partition plant interactions from the pure environment driven plant co-occurrence signal. Our results show that plant interactions shifted from predominantly positive interactions (e.g. nurse-plant facilitation) during the glacial period to negative interactions (e.g. competition) during the Holocene. This shift coincided with a community transition from herbaceous to woody taxa, leading to an increase in average plant height and root length, as evidenced by leveraging plant trait information. We suggest that climate (an external factor) and plant interactions (an internal process) jointly supported rapid and widespread woody taxa expansion at the end of the last glacial, which may provide an analogy with contemporary “arctic greening”. In turn, woody encroachment is likely to constrain the geographical ranges of native species, increasing the risk of local native taxa loss, while enhancing beta diversity.

## Introduction

Woody taxa expansion and increased biomass have been widely observed across the circumpolar region, which is referred to as Arctic greening ^1–4^. This process of woody encroachment has been reported to reduce the diversity and cover of other functional groups ^5,6^. However, certain shrub species also act as nurse plants, exerting positive interactions that facilitate the establishment and persistence of neighbouring taxa ^7^. Yet, all the understanding of these ecological dynamics remains constrained by short-term observations spanning only a few decades, leaving critical uncertainties regarding the long-term consequences of woody plant expansion. How will plant community turnover respond to Arctic warming? How will plant interactions evolve under sustained woody encroachment? What factors drive these shifts, and how do they, in turn, shape the plant community? Addressing these questions requires a long-term perspective and a deeper understanding of the mechanisms governing plant interactions in a changing Arctic.

According to the Stress Gradient Hypothesis, positive interactions among plants are more prevalent in stressful environments, whereas negative interactions dominate under more favourable conditions ^8^. An empirical example of this hypothesis is cushion plants—low-growing, mat-forming genera such as *Eritrichium*, *Saxifraga*, and *Draba*—which survive in harsh Arctic and alpine environments ^9–11^. These plants enhance local microhabitats by trapping heat, stabilising soil, and facilitating the establishment of other species within their crown areas ^12,13^. While cushion-plant studies provide insights, empirical investigations of other plant interactions in Arctic ecosystems remain limited, constraining our capacity to predict future shifts in community dynamics under ongoing environmental changes.

Plants interact with each other through phenotypic characteristics, which reflect strategies of resource acquisition and allocation, mediate interactions with neighbouring plants, and ultimately shape plant–plant dynamics ^14^. Plant height is a key determinant in pre-empting light resources and enhancing competitive ability. Across much of the Arctic, plant height has been increasing in response to warming ^15^, potentially intensifying competitive interactions. Similarly, root length, a critical trait governing belowground water and nutrient uptake ^16^, may further drive competition as deep-rooted taxa expand. Different plant growth forms vary not only in plant height, but also in root length, which can shape the plant interactions. However, how woody encroachment corresponds to community level traits, and how these resulting shifts in traits shape community-wide interactions, are largely unknown. This highlights the need for further investigation into the connections between plant traits and interaction dynamics.

The shifts in plant communities and interactions from the glacial period to the Holocene, driven by rising temperatures, provide a past analogue for contemporary ecological change in the Arctic. Northeast Siberia and Alaska, in particular, have experienced pronounced temperature shifts over millennial timescales due to polar amplification. Furthermore, this region was largely free from ice sheets during the last glacial period ^17,18^, minimising lagged species assembly effects as new habitats formed. High-resolution lake sediment archives, extending back to the Last Glacial Maximum, combined with sedimentary ancient DNA (sedaDNA) records, provide robust datasets for reconstructing plant community dynamics ^19^. Such datasets have enabled the reconstruction of shifts in the relationship between species richness and mean range size ^20^, as well as patterns of taxa loss ^21^. The higher taxonomic resolution of sedaDNA ^22–24^, compared to pollen analysis, facilitates a more accurate reconstruction of taxon interactions through integrative network analysis ^25,26^. Mathematical approaches provide an efficient means of disentangling the mechanisms underlying plant co-occurrence, thereby enhancing our understanding of species assembly and community dynamics. Simple null models, which employ randomised species occurrence matrices, attribute deviations from randomness to species interactions, although they fail to account for the effects of climate and dispersal limitation ^27^. In contrast, environmentally constrained null models partially control for environmental influences while incorporating species’ geographical distributions to account for dispersal effects ^28^. This approach has not yet been applied to investigate long-term species interactions using proxy data.

In this study, we leverage sedaDNA-based plant community reconstructions to investigate plant interactions over time, using nine lake sediment records from northeast Siberia and Alaska spanning the last 24,000 years. To disentangle the effects of plant interactions from other factors shaping reconstructed plant co-occurrence patterns, we apply environmentally constrained null models. Additionally, we examine key functional traits, including plant height and maximum root length, which influence light competition and resource acquisition and may consequently shape species interactions. Our study aims to (1) quantify changes in plant growth-form and functional traits associated with postglacial woody taxa expansion, (2) explore the dynamics of plant interactions throughout this process, and (3) identify the drivers of plant interaction shifts and their role in shaping plant communities.

## Results

### Postglacial woody taxa expansion in response to warming

The study area covers northeastern Siberia and Alaska (Figure 1a). We reanalysed a sedaDNA dataset originally generated by Courtin et al. ^21^ and Jia et al. ^29^, incorporating an additional site, Salmon Lake. All data use metabarcoding approaches targeting the P6 loop of the *trn*L (UAA) intron. Following data filtering ^21,29^, only samples originating in the past 24,000 years and amplicon sequence variants (ASVs) with 100% identity to the customised regional SibAla_2023 database ^21^ were retained. The final dataset consists of 369 samples, generating a total of 78,088,278 reads, representing 625 ASVs.

**Figure 1.**
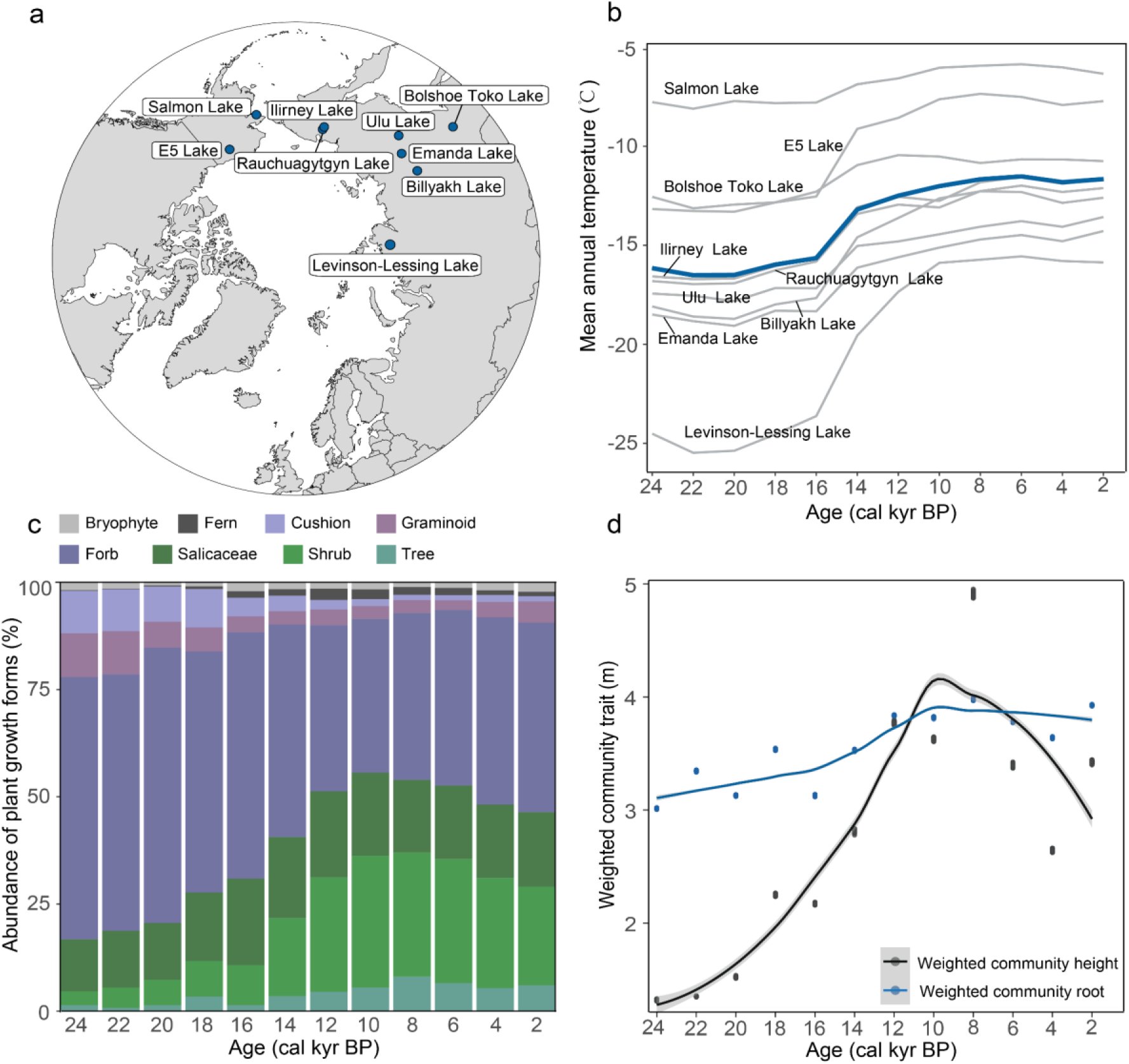
Mean annual temperature, plant growth-form abundance, and community weighted trait over time. (a) Locations of the nine lakes in northeastern Siberia and Alaska. (b) Temporal variations in mean annual temperature (see Materials and Methods). The thin grey curves show site-specific mean annual temperatures, whereas the bold blue curve represents the mean across all study sites. (c) Temporal dynamics in the proportion of transformed counts, representing the relative abundance of distinct plant growth-forms, are depicted in different colours. (d) Temporal changes in community-weighted traits. Traits were derived by summing the trait values of individual ASVs weighted by their relative abundances.

We divided the timeline into 12 time slices, each spanning 2,000 years. For each time slice, the mean annual temperature at each study site was downscaled based on Kleinen et al. (2023) ^30^ (see Materials and Methods). The results indicate that temperatures in the study region remained relatively low during the glacial period (before 14 cal ka BP), increased during the transition phase, and reached relatively high levels during the Holocene (after 10 cal ka BP) (Figure 1b). Spatially, temperature shows a clear decreasing trend from the Alaskan lake region toward the Siberian lake region (Figure 1b).

Relative abundance data indicate a woody taxa encroachment into the plant community from the glacial period to the Holocene (Figure 1c; Supplementary Figure 1). The community was dominated by herbaceous taxa prior to 14.0 cal ka BP, with forbs comprising approximately 60% of the flora, while cushion plants and graminoids each accounted for less than 10% (Figure 1c). In contrast, woody taxa were comparatively sparse, with a total abundance below 25%. After 14.0 cal ka BP, woody taxa increased in abundance, with Salicaceae rising moderately from 15% to ∼20%. Concurrently, other shrub taxa expanded rapidly, peaking at 29% at the onset of the Holocene (∼10.0 cal ka BP). Tree taxa also show a slight increase, reaching approximately 5%. In contrast, cushion plants and graminoids declined gradually, both falling below 5%. Forbs underwent a marked decline, reaching a minimum of 34% around 10.0 cal ka BP, before gradually recovering.

Two community-level plant functional traits associated with competitive ability—light acquisition (plant height) and resource absorption (maximum root length)—were inferred from sedaDNA-derived data. The results indicate that community-weighted plant height increased between ∼16 and 14 cal ka BP, reaching nearly 3 m. It continued rising and peaked at approximately 5 m during the early Holocene (10–8 cal ka BP), followed by a slight decline while remaining above glacial levels (Figure 1d). In parallel, community-weighted root length began increasing around 14 cal ka BP, reached its peak in the early Holocene (10–8 cal ka BP), which is around 0.8m, and remained relatively high thereafter (Figure 1d).

### Environmentally constrained plant interaction network

We used species distribution models (SDMs) to predict plant occurrence patterns driven by temperature, incorporating a random effect to account for stochastic variations beyond temperature influences. The C-score, derived from species occurrences in sedaDNA data, was compared with that obtained from environmentally constrained null models (SDM-constrained). The plant co-occurrence patterns from sedaDNA with a C-score significantly different from the null model were retained (see Materials and Methods for details). To account for the potentially confounding effect of dispersal limitation, we examined spatial range overlap between species pairs. Those with disjunct or partially overlapping distributions were interpreted as being constrained by dispersal. By systematically ruling out temperature filtering, dispersal limitation, and stochasticity, we attribute the residual co-occurrence patterns to direct plant–plant interactions, providing insights into their temporal dynamics. The overlap between the inferred plant interactions and the Globi plant interaction database is approximately 26.0%, which is significantly greater than that observed in random plant interactions.

Across all time slices, cushion plants exhibit a higher proportion of positive interactions, with a median of ∼60% positive to ∼40% negative interactions. In contrast, woody taxa, including trees and shrubs, are predominantly associated with negative interactions, comprising 70–95% of their total links. From the perspective of taxa composition within the plant–plant interaction network, forbs contribute the highest number of species, which are involved in both positive and negative interactions. (Figure 2a, Figure 2d, Figure 2e). Among other herbaceous groups, cushion plants and graminoids are predominantly engaged in positive interactions, particularly during the glacial period (24.0–14.0 cal ka BP). In contrast, woody taxa participate in both positive and negative interactions, with a marked shift toward predominantly negative interactions during the Holocene period (after 14.0 cal ka BP), for example, *Larix*.

**Figure 2.**
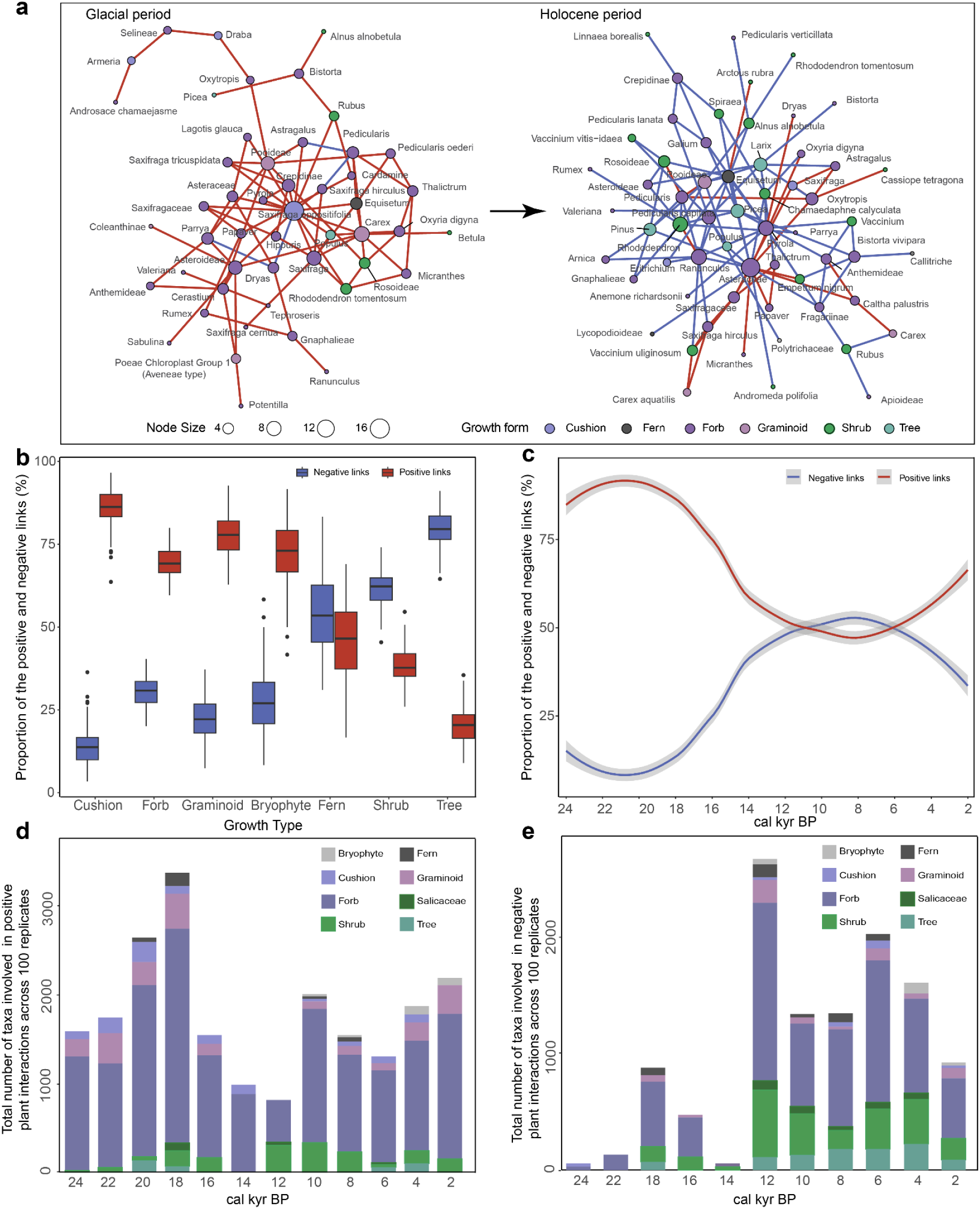
Dynamic plant interaction network during the glacial and Holocene periods. (a) The left panel shows a representative network from 20–18 cal ka BP, illustrating the pattern of the glacial period, while the right panel displays a network from 6–4 cal ka BP, representing the Holocene pattern. Nodes represent taxa, with different growth-forms indicated by colour. Node size corresponds to the number of connections (i.e., node degree). (b) Boxplot of positive and negative interaction percentages across different growth-form taxa. (c) Temporal trends in the relative proportion of positive and negative interactions. (d) Temporal variation in the number of taxa with different growth-forms engaged in positive interactions. (e) Temporal variation in the number of taxa with different growth-forms involved in negative interactions.

From the perspective of interaction direction, positive plant interactions dominated during the glacial period, peaking around the Last Glacial Maximum (20–18 cal ka BP), with broad taxonomic involvement, for example, *Saxifraga oppositifolia* (Figure 2a, 2d, Supplementary Figure 2). By ∼14.0 cal ka BP, negative interactions began to increase (Figure 2a, 2e), reaching equilibrium with positive interactions at the onset of the Holocene (ca. 10 cal ka BP). Thereafter, negative interactions became predominant but decreased again during the Late Holocene (Figure 2c, Supplementary Figure 2).

### Potential drivers of the negative link proportion

A structural equation model was used to assess the direct and indirect effects of temperature and plant community functional traits on the proportion of negative plant interactions. Because of strong collinearity with community mean height and root length (Figure 3a), woody plant abundance was excluded from the final model to improve parameter stability and model fit. The model has a good fit to the data, as indicated by the fit metrics: chi squared (χ²) = 0.3, degrees of freedom (df) = 1, p = 0.6, Root Mean Square Error of Approximation (RMSEA) = 0.0, Comparative Fit Index (CFI) = 1.0, Goodness of Fit Index (GFI) = 1.0, and Standardised Root Mean Square Residual (SRMR) = 0.001. The results reveal that community plant traits responded strongly to increasing temperatures, with plants growing taller and developing longer roots, which in turn led to more negative plant interactions. All effects were statistically significant (p<0.05), as assessed by Wald z-tests based on maximum likelihood estimation.

**Figure 3.**
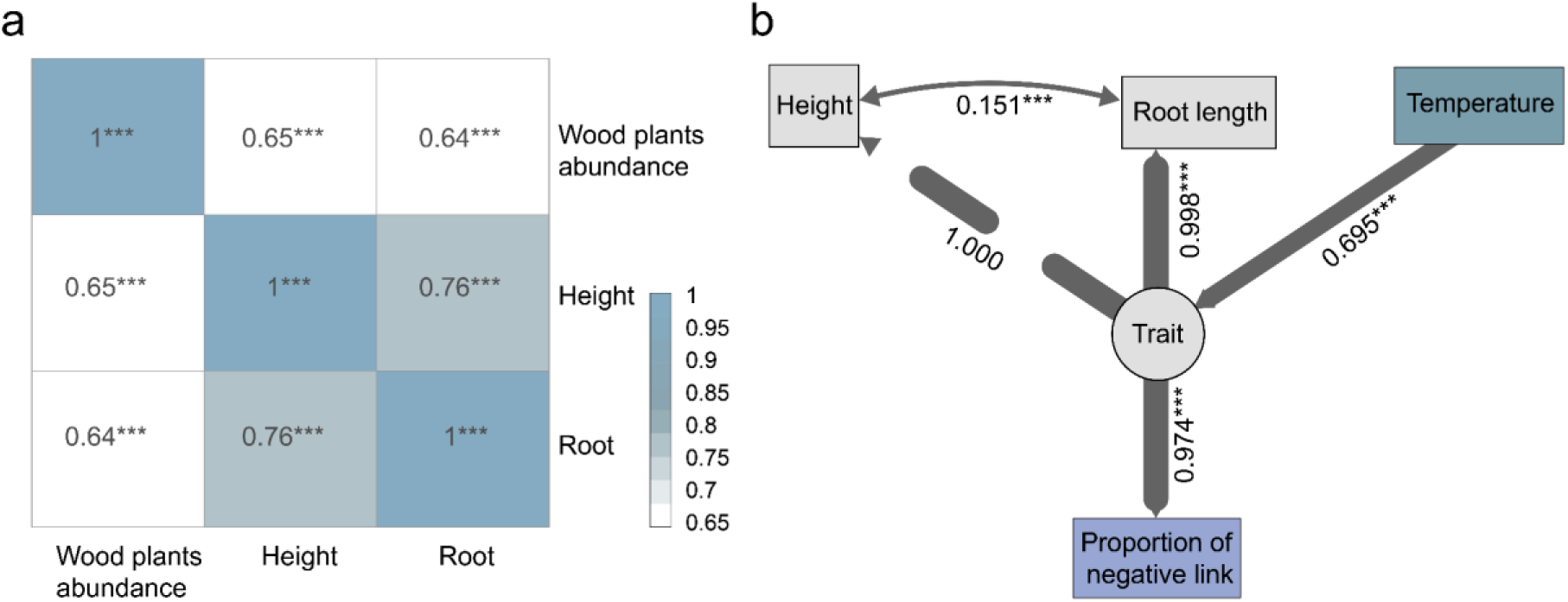
Structural equation model showing the relationships between response variables and predictors. (a) Correlation matrix between woody taxa abundance and two plant functional traits. (b) The structural equation model illustrates the effects of temperature and plant traits on the proportion of negative links. Variables are colour-coded by origin: dark blue for temperature, light grey for plant functional traits, and light bluish purple for the proportion of negative interactions. ‘Trait’ was modelled as a latent variable, inferred from two observed indicators: height and root length. Arrows originate from the latent construct ‘Trait’ and point toward its indicators, height and root length, representing factor loadings. The factor loading of height on the latent construct was fixed to 1, indicated by a dashed arrow. Double-headed arrows represent covariances between observed variables, single-headed arrows indicate regression paths. Arrow thickness reflects the magnitude of the standardised regression coefficients or covariance. Standardised path coefficients and covariances are displayed alongside the corresponding paths. *Significant at the 0.05 level; **significant at the 0.01 level; ***significant at the 0.001 level. Model fit was assessed using multiple metrics: a relatively small chi-square value (χ²), a non-significant p-value (p > 0.05), RMSEA < 0.05, CFI > 0.95, GFI > 0.95, and SRMR ≤ 0.05 collectively indicate a good model fit.

## Discussion

### Plant interactions inferred from the sedaDNA-based co-occurrence networks

We find that herbaceous plants are involved in more positive plant interactions, whereas woody taxa are associated with more negative interactions across all time slices. The observed plant interactions across different growth-forms align with modern ecological expectations. Cushion plants, known to act as nurse species in harsh environments, exhibit predominantly positive interactions with neighbouring plants, such as facilitation and cooperation ^9,13,31^. Consistently, our co-occurrence analysis reveals that cushion plant taxa form a higher proportion of positive than negative links. In contrast, woody taxa, particularly trees, are more frequently associated with negative interactions, consistent with reports of their competitive interactions with neighbouring taxa ^6,32^. The consistent interaction patterns across different growth-forms further support the robustness of the identified plant interaction network.

We inferred reasonable plant interactions from environmentally constrained co-occurrence networks following protocols applied to modern plant co-occurrence, such as those sampled along a broad elevational gradient in the Swiss Alps, but new to palaeoecological studies. This analysis incorporates the potential influences of climate, dispersal limitations, and stochastic processes ^28^. Furthermore, the overlap between the inferred plant interactions and the Globi plant interaction database is significantly greater than that observed in random plant interactions, thereby supporting the reliability of the inferred plant interactions.

Our analysis, however, still has certain limitations. The climatic effects are filtered through a species distribution model (SDM), which inherently introduces errors, such as insufficient species sampling sites, potentially retaining false links or omitting true ones. Additionally, pairwise calculations may overestimate associations driven by covariation, which can vanish when additional species interactions are considered ^33^, potentially allowing indirect interactions to be misinterpreted as direct. However, this method mitigates the risk of overlooking interactions, where some associations are over-interpreted through partial correlations. Also, negative interactions may be underestimated, as strong competition can prevent co-occurrence or drive species exclusion. Furthermore, our study infers plant interactions using DNA samples, with a maximum of 18 samples per 2,000-year time slice. To enhance the robustness and resolution of network inferences larger sample sizes are required ^34^. The taphonomy processes affecting DNA preservation in lake sediments may also bias the representation of the original plant community ^35,36^. Although the use of presence-absence data mitigates some of these biases, it does not fully eliminate them. Lastly, the resolution limitations of amplicon sequence variants (ASVs) prevent the precise taxonomic identification of all taxa to the species level. In the resampled dataset, 55.2% of ASVs were identified at species level, 31.2% at the genus level, and 13.6% at the family level. A single ASV may correspond to multiple taxonomically related species but is represented by only one interaction pair, potentially leading to an underestimation of interaction links.

### Effects of woody encroachment on plant interaction shifts

The proportion of negative plant taxa interactions increases with woody taxa encroachment. During the glacial period (before 14 cal ka BP), the study region remained largely ice-free ^17,37^, providing the required habitat for species survival. Simulations and proxy-based temperature reconstructions suggest relatively low temperatures during this period ^30,38^. Harsh climate conditions likely acted as an environmental filter, favouring the persistence of cold-adapted herbaceous taxa, including forbs, cushion plants, and graminoids. The survival of woody taxa was restricted to specific refugial sites with favourable microclimatic conditions ^29,39–41^. The plant interactions were mainly positive, with most taxa being herbaceous. As the climate warmed from the glacial period to the Bølling–Allerød Interstadial (14–12 cal ka BP), the study area became increasingly favourable for relatively thermophilic species (Figure 1b), promoting the expansion of woody taxa. Key taxa contributing to this process include willow (*Salix* spp.), alder (*Alnus* spp.), birch (*Betula* spp.), and *Vaccinium* spp. With woody taxa encroachment, negative plant interactions increase, involving both woody and herbaceous species.

The higher height of woody taxa may contribute to aboveground light competition when they encroach. Plants interact through their phenotypic characteristics, and changes in plant functional traits can drive shifts in these interactions ^42^. Erect shrubs and trees, such as *Alnus*, *Betula*, *Salix*, *Picea*, *Abies*, and *Larix*, are on average taller than herbaceous plants ^43^, which, when coupled with relatively favourable conditions, enable them to achieve greater heights. Their encroachment contributed to an overall increase in plant height during the Bølling– Allerød Interstadial. This, in turn, raised the community-weighted plant height, with woody taxa intensifying light competition and reducing light availability for low-lying plants ^44,45^. Such changes can lead to etiolation (the process by which plants exhibit elongated growth, pale colouration, and poor health) or exclusion, particularly for shade-intolerant species ^46^. The plant height peaked in the early Holocene, which was followed by a slight decline due to the expansion of dwarf shrubs.

The expansion of woody taxa, characterised by their relatively long root systems, likely enhanced belowground competition for nutrients and space. Root system architecture is a key functional trait governing water and nutrient acquisition ^16^. Coinciding with the expansion of woody vegetation, relatively long root length likely intensified belowground competition for resources and space, for example, for *Picea* and *Pinus* ^43,47^. Additionally, the spread of woody taxa facilitated the formation of mycorrhizal associations ^48^, further enhancing nutrient and water uptake. For example, ectomycorrhizal fungi are commonly associated with the roots of Betulaceae, Salicaceae, and Rosaceae, while ericoid mycorrhizas form symbiotic relationships with members of Ericaceae, such as *Vaccinium vitis-idaea* and *Cassiope tetragona* ^49^. Consequently, community-level root length increased during the Bølling–Allerød interstadial and remained elevated throughout the Holocene, suggesting intensified belowground competition for resources within plant communities.

The structural equation model identifies key drivers of negative plant interactions (Figure 3). As temperatures rise, the increasing abundance of woody taxa enhances community canopy height and root length, intensifying competitive interactions. While temperature indirectly influences these dynamics, shifts in functional traits associated with woody expansion serve as the direct driver of increased negative interactions.

Some woody taxa shift from acting as nurse plants, which primarily interact positively, to becoming competitors with woody taxa encroachment. Plant interaction networks suggest that positive interactions are largely confined to forbs and other herbaceous taxa. Certain woody species also contributed to facilitation and cooperation within plant communities during glacial periods, in contrast to the predominantly negative interactions of woody taxa during interglacial periods. Among them, some shrubs have been documented as nurse plants, alleviating extreme microenvironmental stresses such as drought, elevated temperatures, and soil surface instability ^50^. For instance, the dwarf shrubs *Cassiope tetragona* and *Empetrum nigrum* have been shown to provide aboveground shelter, as indicated by the increased shoot height of taxa growing in their proximity. Both taxa show mainly positive interactions during the glacial period. This effect is likely driven by enhanced snow cover, protection from strong desiccating winds early in the growing season, and a warmer microclimate ^51^. However, during the Holocene, plant interactions shifted to include negative interactions, likely driven by competition for light and resources, as well as potential disruptions to germination ^57^.

This pattern aligns with the Stress Gradient Hypothesis, which posits that positive interactions dominate under stressful conditions, while competition becomes more prevalent in favourable environments ^8,52^. Such conditional interaction strategies of shrub species may maximise their fitness across varying environments. Facilitative interactions likely enhanced their survival during glacial periods by buffering harsh environmental stresses, while the shift towards competitive interactions in the Holocene may have allowed them to acquire resources in less stressful environments. This ability to switch between facilitation and competition may have supported their persistence across both glacial and interglacial periods.

### Potential effects of plant interactions on the past and future plant community

Our results provide quantitative evidence that woody encroachment intensifies negative plant– plant interactions, thereby influencing plant communities. During the glacial period, when the abundance of woody taxa was relatively low, plant interactions were predominantly positive (Figure 4a). The positive interaction likely facilitated the establishment of new taxa and supported the survival of species that might have otherwise been constrained by environmental conditions ^53^. However, this also rendered the plant community more vulnerable to invasions by other taxa. Positive interactions may have further contributed to the persistence of resident taxa by promoting their geographical range expansion. Our previous study found that, during the glacial period, the average range size of plant communities increased, with species facilitating each other’s distribution ^20^. For instance, *Silene acaulis*, which engages in positive interactions with multiple taxa, played a role in expanding species’ ranges ^9^. As a result, regional beta diversity remained low, as the widespread distribution of species promoted a more homogeneous plant composition. Furthermore, the positive plant interactions and relatively wide distribution range likely prevented the plant from going extinct, sustaining diversity even under harsh environmental conditions ^54,55^.

**Figure 4.**
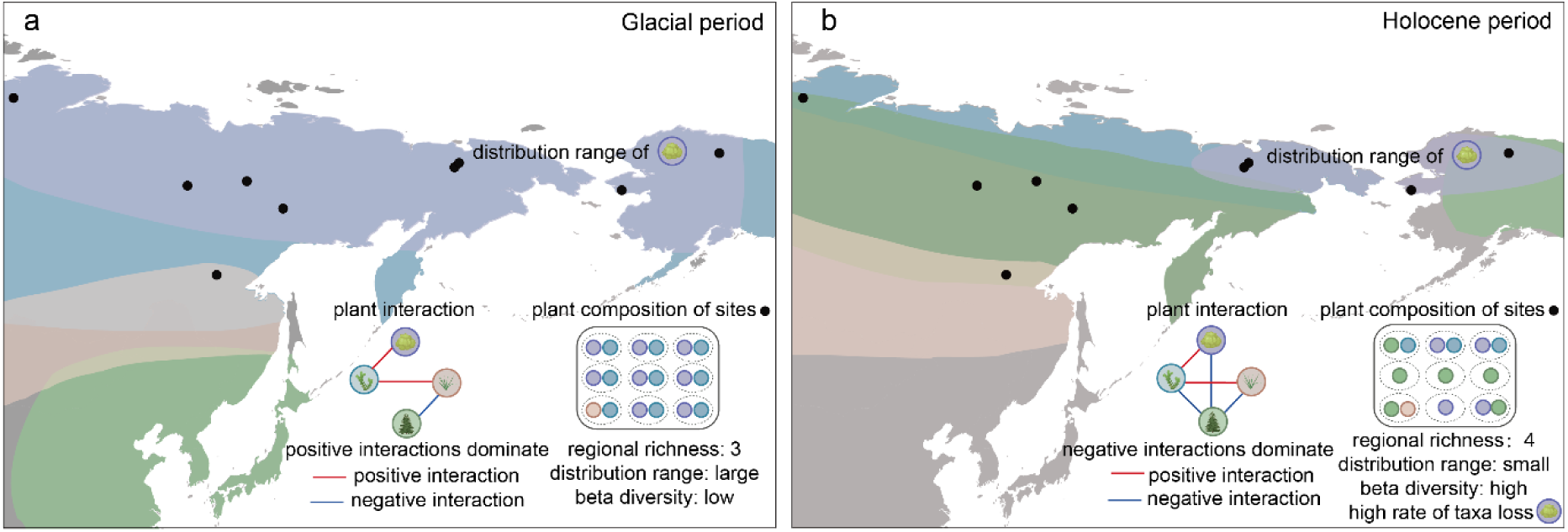
Schematic diagram depicting the impact of plant interactions on community. The coloured polygons represent the distribution ranges of different species, which are presented in the plant interaction network. Red lines indicate positive plant interactions, while blue lines denote negative interactions. Black dots mark the nine sites from which sedaDNA data were obtained, with each site’s plant composition visualised within an ellipse. The combined results from all nine sites form the regional species composition, which is enclosed within a rounded rectangle that contains all the ellipses.

In contrast, the intensified woody encroachment during the interglacial period, which resulted in high woody taxa abundance, increased the incidence of negative plant interactions (Figure 4b). Such intensified negative interactions may constrain the establishment of new taxa by favouring native species with competitive advantages ^56^. As a result, this area exhibits increased resistance to species migration, likely mediated through competition for light and resources, for example for *Larix*, as well as the release of allelopathic chemicals ^57^. Shifts in plant–plant interactions may also influence the persistence of resident taxa, as stronger negative interactions could suppress population growth rates, restricting range expansion and potentially leading to range contraction ^20,58,59^. As a result, these species face an increased likelihood of local extirpation, consistent with the findings of Courtin et al. ^21^, who find a higher than expected loss of taxa, ultimately leading to a decline in diversity. Moreover, herbaceous taxa, due to their traits, are less competitive and particularly vulnerable to local extinction in the study region (Figure 4b, species marked in purple circles), as noted by Courtin et al. ^21^. However, regional beta diversity may increase, as competition fosters divergent community assemblages across sites (Figure 4b), enhancing spatial heterogeneity at broader scales ^60^. Furthermore, our findings suggest that species with advantageous traits under negative interactions are more likely to become dominant in plant communities ^61^ (Figure 4b, woody species marked in green circles).

Overall, while most species distribution models emphasise the direct effects of abiotic factors, biotic interactions such as competition can modify or even counteract the direct impacts of climate on species performance ^62^. Certain plant taxa adjust the direction of their interactions with changing environments. For instance, *Cassiope tetragona* shifts from predominantly positive during the glacial period to primarily negative interactions during the interglacial period. Such conditional interaction strategies may maximise their fitness under the varying environments. This study highlights that woody taxa are likely to dominate Arctic plant communities under climate warming due to their advantageous competitive traits, which facilitate their establishment and expansion. These traits ultimately contribute to their widespread geographical distribution. In contrast, native taxa with lower competitive ability, such as those with shorter stature or shallower root systems, warrant greater attention, as they may decline or disappear before environmental conditions exceed their physiological tolerance, driven by negative interactions. Arctic plant community composition is shaped by more than just the northward migration of lower-latitude species, as biotic interactions may prevent certain herbaceous plants from establishing. Ultimately, plant diversity in the Arctic will be determined by the dynamic balance between species gains and losses.

## Conclusions

Our study used sedaDNA from nine lake sediment cores to reconstruct the dynamics of plant interactions over time in northeast Siberia and Alaska. The results reveal a shift from predominantly positive interactions during the glacial period to increasingly negative interactions in the Holocene. The expansion of woody taxa, driven by rising temperatures and characterised by increased height and root length, contributes to negative interactions such as intensified aboveground light competition and belowground nutrient competition. Certain persistent taxa, such as dwarf shrubs, adjust their strategy accordingly, shifting from positive interactions as nurse plants during glacial periods to negative interactions under changing climatic conditions, which would maximise their fitness. Thus, while temperature acts as an indirect driver, the expansion of woody taxa serves as the direct driver of these negative interactions. Community-level plant functional traits, such as height, may serve as key indicators of plant–plant interactions, offering insights for future conservation efforts. Our findings suggest that future warming in the Arctic will likely lead to a woody taxa-dominated plant composition, intensify negative interactions, and potentially constrain the geographical range of native species, increasing the risk of native taxa loss. However, at a regional scale, more heterogeneous plant composition could enhance beta diversity.

## Materials and Methods

### The sedaDNA dataset

Sedimentary ancient DNA was extracted from nine lake sediment cores collected in northeastern Siberia and Alaska. All sedaDNA processing followed the protocols of a previous study ^21^. In brief, the DNA was isolated from subsamples, purified, and amplified with g and h primers targeting the P6 loop of the chloroplast *trn*L (UAA) intron ^63^. All procedures from extraction to PCR amplification, purifying and merging were conducted in the palaeogenetic laboratories of the Alfred Wegener Institute, Helmholtz Centre for Polar and Marine Research, Potsdam. The purified PCR pools were prepared with a PCR free library and sequenced on the Illumina platform (NextSeq2000 or Nova Seq) in paired-end mode by the company Genesupport Fasteris SA (Switzerland). After DNA sequencing, the raw sequencing files were quality checked and paired-end reads were merged, further sequences were sorted according to samples and taxonomically assigned to the customized SibAla_2023 database ^21^. All steps were performed using OBITools v3 ^64^. Only Amplicon Sequence Variants (ASVs) with 100% matches to the customized “SibAla_2023” database were included in the analysis. Further quality control was conducted based on PCR replicability: PCR samples were excluded if their replicates showed substantial divergence from other samples, indicating low quality, or if replicates failed to cluster, suggesting distinct compositions within replicates ^21^.

The age of each subsample was determined using Bayesian age-depth modelling from previous studies ^29,65–71^, incorporating all samples spanning from 24 cal ka BP to the present. The sedaDNA dataset was divided into 12 time slices, each spanning 2,000 years. To address variations in read counts across lakes and time slices, datasets were rarefied to a base count of 20,000 and resampled 100 times. This base count is lower than the minimum read count observed across all lakes within any given time slice, ensuring the retention of the majority of the original read information.

### Temperature downscaling from the model

Palaeoclimate simulations for mean annual temperature and mean July temperature were obtained from the Max Planck Institute Earth System Model (MPI-ESM1.2), covering the last 23 ka BP ^30,72,73^, with a spatial resolution of approximately 3.75° × 3.75°. The raw temperature data were subsequently downscaled to a resolution of 5 km × 5 km, following the method outlined by Karger et al. ^74^. Briefly, the lapse rate (Γ) was calculated using the average temperature at six pressure levels ranging from 1000 hPa to 600 hPa:

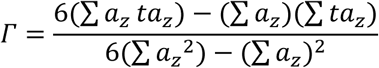

where 𝑡𝑎_𝑧_ is the average temperature at the geopotential height 𝑎_𝑧_and the pressure level Z (6 levels from 1000 hPa to 600 hPa). The palaeo-elevation data was calculated by subtracting past sea level changes ^74^ from modern elevation ^75^.

The downscaled temperature at time slice t (𝑡𝑎𝑠_𝑡_^𝑑𝑠^) was then calculated by:

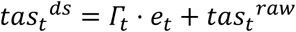

where 𝛤_𝑡_, 𝑒_𝑡_ and 𝑡𝑎𝑠_𝑡_^𝑟𝑎𝑤^ are lapse rate (°C/m), palaeo-elevation (m), and raw temperature (°C) at timeslice t, respectively. Then the downscaled temperature data were cropped to the ‘SibAla’ region (55–90°N, 50–150°E and 40–90°N, 150°E–140°W). To reduce biases in the palaeoclimate simulations, MPI temperatures at 0 ka were adjusted using the modern CHELSA V2.1 climate dataset ^76^. The discrepancies between MPI and CHELSA temperatures at 0 ka were then used to calibrate temperature estimates for other time slices. For areas exposed by sea-level decline during glacial periods (e.g., the Beringia land bridge), temperatures were estimated using K-nearest neighbour interpolation. Ice-sheet cover was excluded from the climate data for all time slices.

### Species distribution model

Species associated with the ASVs detected through sedaDNA data were retrieved from the Global Biodiversity Information Facility (GBIF; accessed on 8 March 2024) for the ‘SibAla’ region (55–90°N, 50–150°E and 40–90°N, 150°E–140°W). Records were filtered by removing (i) unidentified to species level, (ii) fossil specimens, (iii) those with a coordinate resolution <5 km, and (iv) records prior to 1981. As each ASV in the “SibAla_2023” database may correspond to one or more species, the modern distribution of each ASV was determined based on the combined distributions of all associated species and spatially thinned to 15 km resolution, where each 15 km × 15 km grid cell was assigned a binary presence/absence value. If multiple occurrence records were present within a single grid cell, one was randomly selected. ASVs with fewer than 100 occurrences across all grid cells were excluded from modelling due to insufficient sample size.

To define background data for species distribution modelling, we created an inner buffer of 25 km × 25 km and an outer buffer of 200 km × 200 km around each surveyed site. The environment within the inner buffer was assumed to closely resemble that of the surveyed site, facilitating the presence of the target ASV. In contrast, the environment within the outer buffer is considered sufficiently distinct to explain the absence of the ASV in that region. Background points (representing areas where the target ASV was absent) were sampled from the area between the two buffers. Candidate background points were identified based on significant combined differences in mean annual temperature and July temperature relative to the occurrence points of the target ASV, using a one-class support vector machine (OCSVM) classifier ^77^ implemented via the svm function in the R package e1071 ^78^. From these candidate points, a number of background points equal to the occurrence records of the target ASV were randomly selected for model.

Given their important roles in shaping Arctic plant distributions, community composition, and delineating bioclimatic zones ^79^, mean annual temperature, mean July temperature, and mean January temperature, were obtained from the CHELSA v2.1 climate dataset ^80,81^. These data were used to build the species distribution model, which demonstrated that mean July temperature and mean annual temperature were the two primary variables contributing to species distributions. Therefore, these two variables were selected for use in the final model. They were resampled to 5 km × 5 km resolution and assigned to both occurrence and background sites. A MaxEnt model ^82^ was implemented using the R package dismo (v.1.3.14) ^83^ to build the species distribution models, with modern climate data and plant occurrence/absence information used for model training. Parameter tuning was conducted using the ENMevaluate function in the ENMeval package ^84^, which tests regularisation multiplier values ranging from 1.0 to 5.0 in increments of 1 and six different feature combinations (L, LQ, H, LQH, LQHP, LQHPT; where L = linear, Q = quadratic, H = hinge, P = product, and T = threshold) with 5-fold cross-validation, with the best result selected based on the lowest Akaike information criterion (AIC ^85^) value.

Model performance was evaluated using the Boyce index, calculated with the Boyce() function from the modEvA R package ^86,87^. This index compares model predictions with presence/background data. Most ASVs showed high positive Boyce Index values (Supplementary Figure 4), indicating that species occurrences were more frequent than expected by chance in areas of higher predicted suitability, thus reflecting strong concordance between model predictions and observed occurrences. To predict species distributions in the past, downscaled reconstructions of mean annual temperature and mean July temperature were input into the MaxEnt models using the ‘predict’ function. The prediction was repeated five times for each ASV and time slice, and the mean suitability values were calculated. Suitability thresholds were applied using the “10th percentile training presence” criterion, with suitability values below the threshold converted to absence (0). This process produced occurrence suitability values for each ASV in each 5 km × 5 km grid cell across the study region at 1,000-year intervals throughout the past.

### Environmentally constrained null models for inferring plant interactions

SedaDNA data obtained from resampling in each lake and time slice were used to determine species presence/absence information. The procedure follows the protocol of D’Amen et al. ^28^, which disentangles species interactions from environmental filters and dispersal limitation as drivers of species co-occurrence (Supplementary Figure 3). For each species in the 5 km × 5 km grid cell that encompasses the lake, the suitability predicted by the species distribution model (SDM) was used as a proxy for the probability of taxon presence. Thus, as the species distribution model was built based on the mean annual temperature and the mean July temperature, the SDM output reflected temperature-driven occurrence patterns, the sedaDNA data was assumed as the actual occurrence of species.

The strength of association between each species a and b was quantified using the C-score index, calculated as:

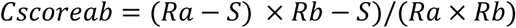

where Ra is the number of sites where species a occurs, Rb is the number of sites where species b occurs, and S is the number of sites in which a and b co-occur. An index value of 0.0 indicates that the species pair is maximally aggregated, with completely nested occurrences. While a value of 1.0 indicates maximal segregation, with the two species never co-occurring. The C-score was calculated using the occurrences of species a and b derived from the sedaDNA data. This value was then compared with the C-score calculated based on the environmentally constrained (SDM-constrained) null models.

For each time slice, the sedaDNA data were arranged such that the rows represent sampling sites and the columns represent species. To generate a null plant community for the same time slice, the species occurrence frequencies (column sums) from the sedaDNA data were preserved, while species richness per plot (row totals) was allowed to vary randomly and equiprobably. Species occurrence sites are determined by the suitability predicted by the SDM, with sites of higher suitability having a greater probability of species occurrence, and vice versa.

The Empirical Bayes method was used to identify statistically significant species pairs. Each pairwise C-score was assigned to one of 22 evenly spaced bins spanning the interval from 0 to 1. The average number of species pairs with different scores in each bin was then calculated from the null communities, representing the null expectation of the C-score for species pairs in that bin. For each bin, the mean limits were computed, species pairs were ordered by their scores within each bin, and those with scores greater than the mean of the simulated distribution (Bayes M criterion) were retained. The set of significant species pairs was further refined by retaining only those that were statistically significant based on an individual test (simple CL criterion). For each species pair, if the C-score derived from the sedaDNA data is significantly higher than that from the environmentally constrained null models, the species are more segregated than expected based on environmental factors. Conversely, if the C-score is lower, the species are more aggregated than expected.

To ensure that species aggregation or segregation is not driven by the geographic configuration of their ranges (e.g., species a and species b being segregated simply due to non-overlapping distribution ranges), the distribution range of each species pair was compared. The Extent of Occurrence (EOO) method ^88^ was used to calculate the distribution range, which defines the minimum convex polygon encompassing the occupied plots. A MANOVA analysis was then conducted to test whether the spatial centroids of the two species’ occurrences differed significantly. If the spatial centroids do not differ, the species pair association can be attributed to positive or negative interactions. If the centroids differ, the association may be due to either interactions or dispersal limitations. In this study, only species pairs with non-differing spatial centroids were considered to represent species interactions.

To assess the reliability of the plant interaction result, the plant interaction pairs were compared with the Globi plant interaction database ^89^ at both the species and genus levels, to determine the proportion of plant interaction pairs that can be found in the database. Additionally, an equal number of interaction pairs were randomly assigned to taxa that were recorded in the sedaDNA dataset but not listed as plant interaction pairs. This random process was repeated 100 times. The overlap between the randomly assigned interaction pairs was compared with the database as well to get the proportion. Finally, the plant interaction overlap rate from the sedaDNA was compared with the rate from the 100-fold randomisations, and the as.randtest() function in the ade4 R package used to test if the overlap rate of the plant interactions is significantly greater than the random rate.

### Plant functional traits

Three functional traits were checked in this study, which are plant growth-form, height, and maximum root length. Growth-form information for each species was primarily obtained from the TRY 5.0 database ^43^; for species lacking database entries, growth-forms were verified manually. For each ASV, the growth-form was assigned based on the most frequent growth-form among the associated species. To account for large differences in read count magnitudes among plant growth-forms, read counts for each ASV were square-root transformed. In each time slice, the relative abundance of each ASV was calculated by dividing its transformed count by the total transformed counts across all ASVs. The abundance of each growth-form was then obtained by summing the relative abundances of all ASVs assigned to that growth-form. Plant height and root length data were obtained from multiple sources, including the Tundra Trait Team (TTT) database ^47^, the TRY 5.0 database ^43^, and the Global Root Trait (GRooT) database ^90^, with additional height data for several species manually retrieved. For ASVs lacking functional trait information, they were not included in subsequent analyses. Functional traits were assigned to each ASV using the mean values of all species corresponding to that ASV. To assess temporal changes of plant functional traits, we weighted each ASV’s functional traits by its relative proportion of counts. The weighted values were then summed to derive community-level functional traits.

### Structural equation model

To detect potential autocorrelation, the correlation between woody plant abundance and plant functional traits was assessed using the “Pearson” method and a “two-sided” alternative with the cor.test function from the stats package in R. Based on the correlation analysis, we constructed a structural equation model (SEM) according to a conceptual framework, incorporating community mean plant height, root length, and mean annual temperature as observed variables. Negative plant interaction was specified as the response variable. The ‘Trait’ was modelled as a latent variable—an unobserved factor inferred from two observed indicators: height and root length. As latent variables lack inherent measurement scales, the factor loading of height on the latent construct was fixed at 1 to define the scale of the latent dimension. In the theoretical framework, traits were assumed to be influenced by temperature and, in turn, to mediate negative interactions among plants. To evaluate this conceptual framework, we specified a structural equation model (SEM) using the sem() function from the lavaan package in R and assessed its overall goodness-of-fit. All statistical analyses were conducted in R (version 4.3.1) ^91^.

## Data Availability

The raw sedaDNA sequence data have been deposited in the European Nucleotide Archive (ENA) at EMBL-EBI under accession number PRJEB76237 (https://www.ebi.ac.uk/ena/browser/view/PRJEB76237) and under accession number PRJEB70434 (https://www.ebi.ac.uk/ena/browser/view/PRJEB70434). The data generated in this study and used for analyses are provided in the Source Data file.

## Code Availability

R scripts for processing the data, with dataset input provided with this paper, can be downloaded via Zenodo.

## Acknowledgements

We thank Cathy Jenks at the University of Bergen for proofreading this manuscript. This research has been supported by the European Research Council (ERC Glacial Legacy 772852 to Ulrike Herzschuh), and China Scholarship Council (grant 202106620011 to Ying Liu).

## Author Contributions Statement

**Ying Liu**: Conceptualisation, Methodology, Data curation, Software, Formal analysis, Investigation, Visualization, Writing – original draft, Writing – review & editing; **Weihan Jia**: Methodology, Writing – review & editing; **Sisi Liu**: Methodology, Writing – review & editing; **Simeon Lisovski**: Methodology, Writing – review & editing; **Kathleen R. Stoof-Leichsenring**: Methodology, Data curation, Writing – review & editing; **Luca Zsofia Farkas**: Investigation, Writing – review & editing; **Ulrike Herzschuh**: Conceptualisation, Methodology, Resources, Writing – review & editing, Supervision, Funding acquisition.

